# Non-cytolytic re-engineering of a viral vaccine vector enables durable effector-memory T cell immunity by reinforcing type I IFN induction

**DOI:** 10.1101/2025.11.04.685127

**Authors:** Matias Ciancaglini, Robin Avanthay, Anna-Friederike Marx, Tiago Abreu-Mota, Davide Finozzi, Jonas Fixemer, Florian Geier, Dominik Burri, Ingrid Wagner, Ilena Vincenti, Mario Kreuzfeldt, Doron Merkler, Gert Zimmer, Daniel D. Pinschewer

**Affiliations:** Department of Biomedicine, Division of Experimental Virology, University of Basel, 4009 Basel, Switzerland; Institute of Virology and Immunology, 3147 Mittelhäusern, Switzerland and Department of Infectious Diseases and Pathobiology, Vetsuisse Faculty, University of Bern, 3012 Bern, Switzerland; Department of Biomedicine, Bioinformatics Core Facility, University of Basel, 4031 Basel, Switzerland and Swiss Institute of Bioinformatics, Basel, Switzerland; Department of Pathology and Immunology, University of Geneva, 1211 Geneva, Switzerland; Division of Clinical Pathology, Geneva University Hospital, 1206, Geneva, Switzerland

## Abstract

Replication-deficient viral vector systems hold promise for CD8 T cell-based vaccination, but the molecular mechanisms accounting for platform-specific differences in immunogenicity remain ill-defined. When comparing prototypic single-cycle vaccine delivery platforms we found that lymphocytic choriomeningitis virus-based vectors (rLCMV), which are non-cytolytic, elicited more durable and effector-memory-differentiated CD8 T cell responses than vectors based on cytolytic vesicular stomatitis virus (rVSV). Hence we re-engineered rVSV to be non-cytolytic (rVSVMq). This vector induced more durable and effector-differentiated CD8 T cell memory than the parental rVSV and it afforded superior protection against Listeria challenge. Improved CD8 T cell responses of non-cytolytic rVSVMq were driven by a reinforced type I interferon (IFN-I) response and its direct sensing by vaccination-induced CD8 T cells. Many vector cargo-specific CD8 T cells in the splenic marginal zone of rVSVMq- or rLCMV-vaccinated mice were in contact with vector cargo-expressing cells that co-expressed type I interferon. In contrast, rVSV-vectored cargo-expressing contacts of specific CD8 T cells were largely IFN type I-negative. Thereby, vaccination with non-cytolytic viral vectors offered an opportunity for CD8 T cells to integrate peptide-MHC and IFN-I signals during priming. These mechanistic insights should help to refine vaccines aimed at eliciting durable and protective effector-memory CD8 T cell immunity.

## Introduction

CD8 T cells represent a key pillar of adaptive immunity and an important line of defense against intracellular bacteria, tumors and most viral diseases^1–7^. The recent severe acute respiratory syndrome coronavirus 2 (SARS-CoV-2) pandemic resulted in the rapid clinical development of vaccines that were based on novel platforms and has provided additional insights into the role of CD8 T cells in infection and vaccination. CD8 T cells represent a critical component of immunity against severe COVID-19 disease^8^ and are indispensable to achieve control of SARS-CoV-2 replication^9^. Moreover, vaccination-induced CD8 T cell responses are more durable than antibody titers^10^ and they can protect against severe SARS-CoV-2 infection in the absence of neutralizing antibodies^11,12,13^. Equally importantly, rapidly evolving SARS-CoV-2 readily escapes antibody responses^14^ but remains susceptible to recognition by vaccination-induced CD8 T cells, which typically cross-react to a wide range of viral variants^13,15,16^. Under certain conditions CD8 T cell immunity may even prevent the transmission of a respiratory virus^17^.

The development of CD8 T cell-based vaccines has long represented a difficult task, which in recent years has been eased by the advent of new delivery technologies including a variety of viral vectors^18,19,20^. Replication-competent viral vector platforms can, however, raise safety concerns ^21,22^, especially in the context of prophylactic vaccination. These considerations argue in favor of replication-deficient vectors, some of which can induce immune responses that in magnitude are comparable to their replication-competent counterparts ^23,24,25^. Vast differences can, however, be observed between viral vector platforms in their ability to elicit long-lived protective CD8 T cell immunity^19,26^, and it remains insufficiently understood which molecular determinants of viral vectors account for this important feature. Viral targeting to and activation of antigen-presenting cells is widely seen as one important mechanism^19,27–29^. Moreover, the amount and duration of antigen supply is commonly thought of as a determinant of the potency of CD8 T cell responses^26,30–33^, and in case of replicating viral vaccine vector platforms antigen expression can be augmented by transient blockade of the type I interferon (IFN-I) receptor^34^. In contrast, IFN-I signals have commonly been reported to improve or not to negatively impact the immunogenicity of replication-deficient platforms^26,35,36^. Yet viral vectors differ substantially in the levels of IFN-I they induce^37–40^. A further important characteristic of viral vectors to consider is their cytolytic or non-cytolytic behavior, which can impact the processing of vectorized cargo for presentation on MHC class I (MHC-I)^41^ and viral interactions with the immune system more generally^42^. It remains, however, insufficiently understood how IFN-I induction and cytolytic behavior of replication-deficient viral vector platforms are interconnected and how these features translate into durable effector-memory CD8 T cells induction.

Vesicular stomatitis virus (VSV), the prototype member of the rhabdovirus family, has been studied by generations of immunologists for its ability to elicit potent and long-lived humoral immunity^43–46^. A VSV-based vaccine vector (rVSV/ZEBOV) expressing the Zaire Ebola virus (ZEBOV) glycoprotein has been tested in the 2014-2015 Ebola virus epidemic, demonstrating 100% prophylactic efficacy against Ebola virus disease^47–49^, and analogously engineered vaccines against Lassa fever are currently undergoing clinical development^50^. rVSV/ZEBOV induces T cell responses, too, but studies in NHPs showed that protection against lethal challenge infection required vaccination-induced antibody immunity^48,51^. In stark contrast, an adenoviral vaccine expressing the same antigen protected NHPs by inducing CD8 T cell immunity^4^, suggesting that the two platforms differ in the arms of adaptive immunity they preferentially engage. One distinguishing feature of VSV and derived vector system consists in a prominent host cell shut-off, which is mediated by the viral matrix protein M that specifically blocks the nuclear export of cellular RNAs to the cytosol^52–54^. The host shut-off does not affect viral replication and transcription in the cytosol, ascertaining high-level viral protein expression, but it results in rapid cytolytic cell death^55,56^, forming a basis for the utility of VSV and related rhabdoviruses in oncolytic cancer therapy^57,58^. Importantly, the M protein-mediated host shut-off is the only mean how this virus supresses the synthesis and release of type I interferon^59^. Whether and how a cytolytic life with host cell shut-off impacts CD8 T cell induction remains untested.

Here we show that rVSV, when engineered to replicate in a non-cytolytic manner (rVSVMq), elicits more durable and effector-differentiated CD8 T cell memory with improved protective capacity. Superior immunogenicity of rVSVMq depended on its ability to elicit higher levels of systemic IFN-I, which signaled directly to antigen-specific CD8 T cells. Our observations provide a mechanistic link between non-cytolytic replication, systemic IFN-I responses and potent effector-memory CD8 T cell immunity, which should help to rationally refine vaccination strategies.

## Results

### rLCMV induces more durable effector-memory CD8 T cell responses than rVSV

To study and compare the ability of rVSV and rLCMV vectors to induce CD8 T cell immunity, we employed replication-deficient vectors wherein the glycoprotein gene was replaced by the prototypic model antigen ovalbumin (OVA; rLCMV-OVA, rVSV-OVA). Both vectors are genetic vaccine delivery vehicles, requiring intracellular expression of the vector-encoded OVA to elicit immune responses. Accordingly, the production of these glycoprotein-deficient vectors necessitates the use of glycoprotein trans-complementing cell lines, which pseudotype budding particles to become single-round infectious^19,60^. To determine how the choice of viral surface glycoprotein impacted CD8 T cell induction, e.g. by influencing cell tropism, we generated not only rVSV-OVA and rLCMV-OVA carrying their respective own glycoprotein but we produced also rLCMV vectors pseudotyped with VSV glycoprotein (rLCMV/VSVG-OVA) as well as rVSV vectors pseudotyped with LCMV glycoprotein (rVSV/LCMVGP- OVA), all of them single-round infectious. Each one of the four vectors was administered intravenously (i.v.) to mice and OVA-specific CD8 T cell frequencies in blood were measured over time using MHC class I tetramers (Fig. 1A-D, S1A). At day seven after immunization, all vectors elicited similar OVA-specific CD8 T cells (Fig. 1C). By day 20 after prime, however, the responses elicited by rLCMV-OVA and rLCMV/VSVG-OVA remained stable, whereas rVSV-OVA- and rVSV/LCMVGP-OVA-induced CD8 T cells contracted 5- and 4-fold, respectively. These findings suggested that rLCMV-induced CD8 T cell responses were more durable than those induced by rVSV, irrespective of the glycoprotein used to pseudotype the vectors. Not only the durability but also the phenotype of responding CD8 T cells varied greatly between rLCMV- and rVSV-immunized mice. By day 20 after immunization the response to the former consisted predominantly in effector-memory CD8 T cells (KLRG1^+^ CD127^-^) while the response to the latter was dominated by more resting memory phenotype CD8 T cells (KLRG1^-^ CD127^+^) (Fig. 1B,D).

**Figure 1.**
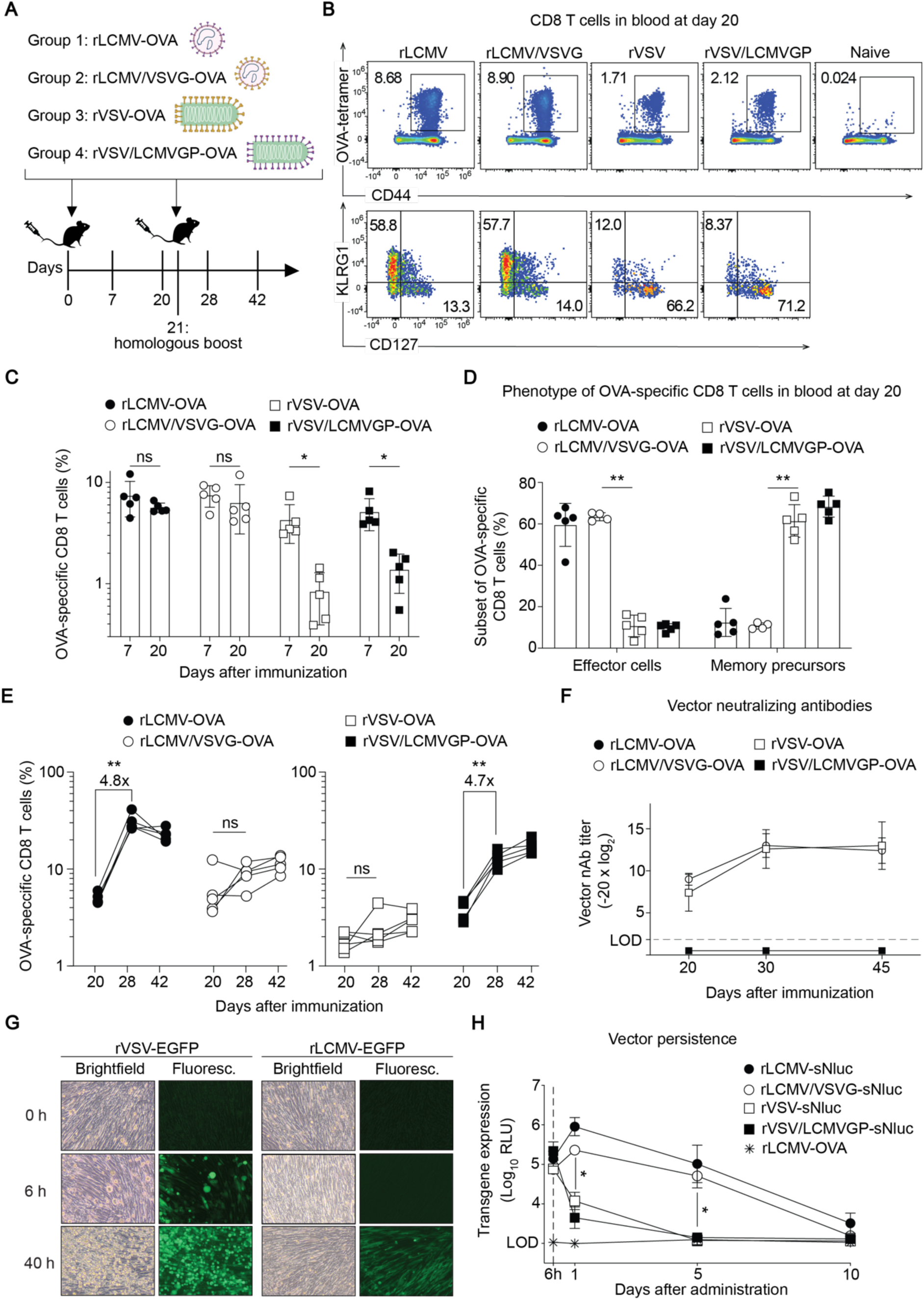
rLCMV induces more durable effector-memory CD8 T cell responses than rVSV. (A) Experimental design. Mice were immunized intravenously with OVA-expressing rLCMV, rVSV or their reciprocal pseudotypes on d0 and d21, and blood as well as serum samples were collected over time. (B) Representative FACS plots of OVA-tetramer-binding CD8 T cells and their phenotype in blood at d20. (C) Frequencies of OVA-tetramer binding CD8 T cells in blood at d7 and d20. (D) Frequencies of OVA-tetramer-binding CD8 T cells with an effector (KLRG1^+^CD127^-^) or memory precursor (KLRG1^-^ CD127^+^) phenotype in blood at d20. (E) Frequencies of OVA-tetramer binding CD8 T cells in blood after primary immunization (d20) and after boost (d28, d42) with rLCMV- or rVSV-based vectors. (F) Vector- neutralizing antibodies in serum of mice vaccinated with rLCMV- or rVSV-based vectors over time. (G) Cytopathic effect of EGFP-expressing rLCMV and rVSV after infection of BHK21 cells. (H) Luciferase activity in serum of WT mice immunized with sNluc-expressing vectors pseudotyped with VSVG or LCMVGP. Samples from mice immunized with rLCMV-OVA were used to determine technical backgrounds. Symbols in (C,D,E) represent individual mice with bars in (C,D) showing the mean±SEM. Symbols in (F) show the mean±SEM of n=5 (F) or n=3 (H) mice. One representative experiment of two similar ones is shown. Statistical analysis was performed by two-way ANOVA with Bonferroni’s post- test for multiple comparisons (C, D, E and H); ns: not significant; *: p < 0.05, **: p < 0.01, *p* > 0.05 was considered not statistically significant and is not indicated.

To test the ability of the various vector formats to boost OVA-specific CD8 T cell response upon re-administration, we performed a second immunization at day 21, in a homologous prime-boost regimen (Fig. 1E). Booster vaccination using vectors pseudotyped with LCMV glycoprotein induced a significant increase in circulating OVA-specific CD8 T cells. By day 28 (seven days after boost), the frequencies of rLCMV-OVA- and rVSV/LCMVGP-OVA-induced CD8 T cells were augmented 4.8 and 4.7-fold, respectively. VSVG-pseudotyped vectors showed a trend towards higher CD8 T cell frequencies after booster vaccination, too, but these differences failed to reach statistical significance. The responses to all four vectors remained stable up to day 42 (21 days after boost). These results showed that even though the viral glycoproteins did not measurably influence the magnitude of vector- induced primary responses, they had a major impact on the secondary response upon homologous boosting, with vectors carrying LCMV-GP allowing for a more efficient boost of antigen-specific CD8 T cell response than VSVG-pseudotyped ones.

Pre-existing anti-vector immunity and notably vector-neutralizing antibodies can interfere with homologous prime-boost immunization regimens^61^, and the glycoproteins of LCMV and VSV differ substantially in their ability to elicit neutralizing antibodies^46^. To determine vector-neutralizing antibody induction by the different vectors, serum samples from immunized mice were assessed for their neutralization activity (Fig. 1F). The sera of mice vaccinated with either one of the two vectors that carried LCMV glycoproteins, i.e. rLCMV-OVA and rVSV/LCMVGP-OVA, were devoid of detectable LCMV-neutralizing antibody activity, both prior to and after homologous boosting. On the contrary, VSV neutralizing antibodies were readily induced after single immunization with VSV glycoprotein-pseudotyped rVSV-OVA and rLCMV/VSVG-OVA, and these neutralizing titers increased further after homologous boost. These data suggested that the differential ability of rVSV and rLCMV vectors to boost CD8 T cell responses were associated with the differential induction of vector-neutralizing antibodies after primary immunization.

Besides differential glycoprotein-mediated cell tropism, an important distinction between VSV and LCMV consists in their cytolytic and non-cytolytic life cycles, respectively. Unlike rLCMV, which replicates without any noticeable cytopathic effect, rVSV causes infected cells to round up and detach within 6 hours after infection (Fig. 1G). Taking into account that these replication-deficient vectors cannot propagate from cell to cell, antigen expression is limited to the first round of infected cells. Henceforth, the cytolytic activity of the vector can represent a limiting factor for the amount of antigen expressed in vivo, which in return is thought of as a major determinant of vector immunogenicity^26,34,62^. In order to quantify vectored antigen expression over time in mice we generated vectors expressing secreted Nano-luciferase (sNluc), which can be sampled from serum to serve as a measure of the total amount of antigen expressed in the animal’s body. Intravenous administration of recombinant sNluc protein revealed that the protein’s *in vivo* half-life ranged below 1 hour, validating serum sNluc activity as a real-time surrogate of protein synthesis by vector-infected cells (Fig. S1B). sNluc-expressing rVSV and rLCMV vectors were pseudotyped with either VSVG or LCMV-GP and were administered to mice i.v. (Fig. 1H). Similar levels of sNluc activity were detected at 6 hours after administration, irrespective of the vector used. By 24 hours after administration, however, sNluc activity in animals receiving rVSV-based vectors had markedly declined, while sNluc expression by rLCMV-based vectors was still on the rise, largely independently of the glycoprotein used for pseudotyping. By day 5, rVSV-expressed sNluc had dropped below detection limits while rLCMV-driven antigen expression subsided by around day 10. These results indicated that rLCMV can better persist in infected cells *in vivo,* expressing antigen for longer periods of time than rVSV. These observations raised the possibility that increased antigen availability for longer periods of time could have provided a better stimulation to CD8 T cells, culminating in the more durable and effector-differentiated CD8 T cell response observed.

### Non-cytolytic rVSVMq induces long-lived effector-differentiated CD8 T cell memory and provides better protection against Listeria challenge than cytolytic rVSV

We hypothesized that abrogating rVSV’s cytolytic activity might improve the CD8 T cell response. For this, we employed an attenuated VSV matrix protein quadruple mutant vector (rVSVMq), which lacks host shut-off activity and does not cause cytopathic effects in cell culture ^63^. In keeping with earlier reports, rVSVMq cytolytic activity was abrogated, whereas rVSV infection induced a pronounced rounding and detachment of Vero E6 cells from the culture flask (Fig. 2A). To test whether its non-cytolytic replication cycle prolonged rVSVMq antigen expression in mice, we turned again to sNluc- expressing constructs (Fig. 2B). By six hours after administration to wildtype mice, rLCMV-sNluc and rVSV-sNluc expressed similar sNluc levels whereas those of rVSVMq-sNluc-inoculated mice were about ten times lower (Fig. 2B, top). In keeping with the results of Fig. 1H, rLCMV-vectored sNluc levels persisted at more or less constant levels up to day three after inoculation and remained over technical backgrounds for more than seven days whereas rVSV-vectored sNluc declined rapidly, reaching technical backgrounds by day 5. Despite the vector’s non-cytolytic behavior, rVSVMq-expressed sNluc activity declined in parallel to the one produced by rVSV, reaching background levels already on day three. The low and transient sNluc expression by non-cytolytic rVSVMq in wildtype mice suggested, therefore, that vector-extrinsic mechanisms influenced its persistence in the infected cell. For instance, interferon responses as well as adaptive immunity can have a major impact on antigen expression levels of viral vectors^34,61,64^. To test the intrinsic ability of the vectors to persist *in vivo* i.e. their persistence in the absence of innate and adaptive host immune responses, we employed mice that lack type I and II interferon receptors as well as T and B lymphocytes (Fig. 2B, bottom). Both non-cytolytic vectors, rLCMV-sNluc and rVSVMq-sNluc, maintained high levels of sNluc expression in these immunodeficient mice for more than 30 days, indicating that these single-round vectors persisted in the cells they had infected. In contrast, antigen levels expressed from the cytolytic rVSV-sNluc showed a steady decline, reaching background levels by around day 15. Given that rVSV and rVSVMq carried the same glycoprotein and hence targeted the same cell types, these data indicated that declining antigen levels observed for rVSV but not for rVSVMq were related to an intrinsic property of the respective vectors rather to the type of cells they transduced. Taken together, these findings suggested that in animals lacking IFN sensing as well as B and T cells, a non-cytolytic life cycle allowed the vector’s persistence at fairly constant levels for prolonged periods of time, whereas cytolytic activity caused progressive loss of antigen expression, which was presumably reflective of host cell death.

**Figure 2.**
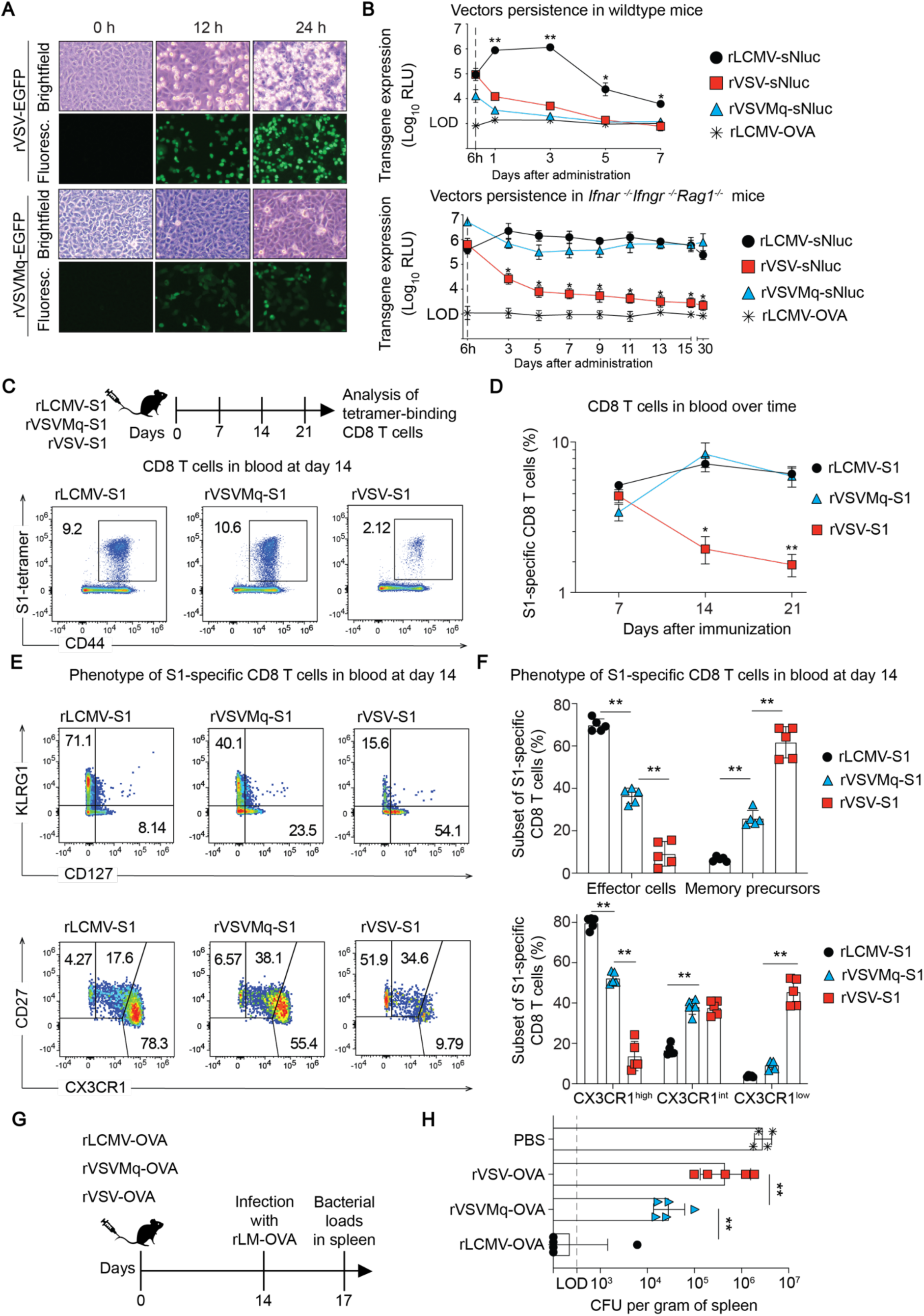
Non-cytolytic rVSVMq induces long-lived effector-differentiated CD8 T cell memory and provides better protection against Listeria challenge than cytolytic rVSV. (A) Cytopathic effect of rVSV-EGFP and rVSVMq-EGFP vectors on Vero E6 cells at 0, 12 or 24 hours after infection. (B) Luciferase activity in serum of WT (top) and *Ifnar^-/-^Ifngr^-/-^Rag^-/-^* mice (bottom) immunized with sNluc- expressing vectors. Samples from mice immunized with rLCMV-OVA were used to determine technical backgrounds. (C) We immunized WT mice with rLCMV-S1, rVSVMq-S1 or rVSV-S1 on d0 and collected blood over time to analyze S1-tetramer-binding CD8 T cells. Representative FACS plots of S1-tetramer- binding CD8 T cells in blood at d14. (D) Frequencies of S1-tetramer-binding CD8 T cells in blood over time. (E) Gating strategy to identify S1-epitope-specific CD8 T cell subsets with an effector or memory precursor phenotype as judged based on KLRG1/CD127 (top) or CD27/CX3CR1 expression (bottom) at d14. (F) Frequencies of effector (KLRG1^+^CD127^-^) and memory precursor (KLRG1^-^CD127^+^) subsets (top) among S1-tetramer-binding CD8 T cells, and the abundance of subsets based on CX3CR1 expression levels in conjunction with CD27 (bottom) at d14 after immunization. (G) We immunized WT mice with rLCMV-OVA, rVSVMq-OVA or rVSV-OVA and 14 days later challenged them with OVA- expressing *Listeria monocytogenes* and collected spleens on d17. (H) Bacterial loads on d17 in spleen. Symbols in (B,D) represent the mean±SEM of n=3 (B) and n=5 (D) mice. Symbols in (F,H) represent individual mice with bars indicating the mean±SEM. One representative experiment of two similar ones is shown. Statistical analyses were performed by two-way ANOVA with Bonferroni’s post-test for multiple comparisons (B, D, F) or one-way ANOVA with Tukey’s post-test (H) *: p < 0.05, **: p < 0.01, *p* > 0.05 was considered not statistically significant and is not indicated.

Next, we set out to corroborate and extend our observations on differential OVA-specific CD8 T cell induction by rVSV and rLCMV (compare Fig. 1A-D) using a different antigen, and to assess the impact of non-cytolytic replication on vector immunogenicity. rLCMV, rVSVMq and rVSV vectors expressing the subunit 1 of the SARS-CoV-2 spike protein (S1) were administered to WT mice and epitope-specific CD8 T cells in blood were analyzed over time using MHC class I tetramers (Fig. 2C,D). Analogously to the observations made with OVA-expressing vectors, the responses elicited by rLCMV- S1 were more durable than those in response to rVSV-S1, which exhibited a more pronounced contraction by day 14 after administration (Fig. 2D, compare Fig. 1C). In remarkable contrast, immunization with rVSVMq-S1 induced frequencies and kinetics of S1 epitope-specific CD8 T cell that were similar to rLCMV-S1. Unlike rVSV-S1-induced CD8 T cell frequencies, which decreased by two weeks after immunization, the rVSVMq response expanded from day seven to day 14 and was maintained at high frequencies until day 21. Similar observations were made in an independent experiment comparing analogous vectors expressing OVA (Fig. S2A,B). Furthermore, we observed analogous kinetics when determining the total numbers of S1-specific CD8 T cells in spleens of mice immunized with the aforementioned vectors (Fig. S2C,D). Another major difference between rLCMV- and rVSV-induced CD8 T cell responses was the phenotype (compare Fig. 1D). rLCMV-S1-induced CD8 T cell responses were dominated by KLRG1^+^CD127^-^ and CX3CR1^+^CD27^-^ effector cells, whereas the rVSV-S1 response was biased towards the KLRG1^-^CD127^+^ and CX3CR1^-^CD27^+^ memory subsets (Fig. 2E,F). In comparison to the latter cytolytic vector, non-cytolytic rVSVMq-S1 induced higher frequencies of both effector subsets. Albeit not reaching the levels obtained after rLCMV immunization, the KLRG1^+^CD127^-^ and CX3CR1^+^CD27^-^ populations were 5- and 4-fold higher than those induced by rVSV-S1, respectively. Similarly, the KLRG1^-^ CD127^+^ and CX3CR1^-^ CD27^+^ memory subsets elicited by rVSVMq-S1 made up for 25% and 10% of the epitope-specific CD8 T cell response, respectively, about midways between rLCMV and rVSV. S1-binding antibody responses on day 7 after rVSV-S1 immunization were significantly higher than those in response to rVSVMq-S1 and a similar trend was noted at later time points (Fig. S2E), indicating that the rVSVMq vector design improved T cell but not B cell immunogenicity.

Finally, we compared the different vaccine vectors in a well-characterized model of CD8 T cell-mediated protection^65^. Mice were immunized with either one of the OVA-expressing vectors and challenged 2 weeks later with OVA-expressing *Listeria monocytogenes* (rLM-OVA), an intracellular bacterium that is controlled by CD8 T cells. Spleens were collected 3 days after infection to determine bacterial loads (Fig. 2G,H). When compared to naïve animals all three vectors conferred some level of protection and mice immunized with rLCMV-OVA were almost free of rLM-OVA. Importantly, rVSV-OVA was only modestly protective while rVSVMq-OVA-induced immunity suppressed bacterial loads to 100- fold lower levels than rVSV-OVA vaccination. Taken together these data showed that the non-cytolytic rVSVMq-OVA induced more durable and more effector-differentiated CD8 T cell memory than rVSV- OVA, which correlated with substantially improved protection against Listeria challenge.

### rVSVMq-induced CD8 T cells exhibit transcriptomic signatures of effector differentiation and IFN-I signaling

The kinetics and phenotype of rVSVMq-induced CD8 T cell responses were dissimilar to those elicited by rVSV but resembled the responses observed in rLCMV-immunized mice, prompting us to characterize them at the transcriptional level. We vaccinated animals with either rVSV-S1, rVSVMq-S1 or rLCMV-S1, sorted tetramer-binding cells seven and fourteen days later and processed them for single cell RNA sequencing (scRNAseq, Figs. 3A, S3A). Projection of the sequencing data onto a t-distributed stochastic neighbor embedding (t-SNE) dimensional reduction space differentiated five clusters of cells (Fig. 3B). The progenitor cluster 1 displayed moderate expression of the memory-associated transcription factor *Tcf7*, the survival cytokine receptor *Il7r*, and the progenitor marker *Slamf6*, in conjunction with intermediate expression of the chemokine receptor *Cxcr3* and moderate expression of effector-related genes such as *Gzmk* (Fig. 3C). The effector cluster 2 contained copious amounts of transcripts for *Gzmb* (cytotoxic activity), the effector-associated surface markers *Cx3cr1* and *Klrg1*, sphingosine-1-phosphate receptor 5 (*S1pr5*) ^66^ and for the effector differentiation-associated gene *Zeb2* ^67^. Early memory cluster 3 cells were rich in transcripts for *Tcf7*, the anti-apoptotic regulator *Bcl2* as well as for the survival receptor *Il7r*. The central memory cluster 4 resembled cluster 3, but exhibited an increased expression of *Sell* (encoding for L-selectin) and CCR7. The interferon signature cluster 5 was enriched for IFN-stimulated gene transcripts such as *Irf7*.

**Figure 3.**
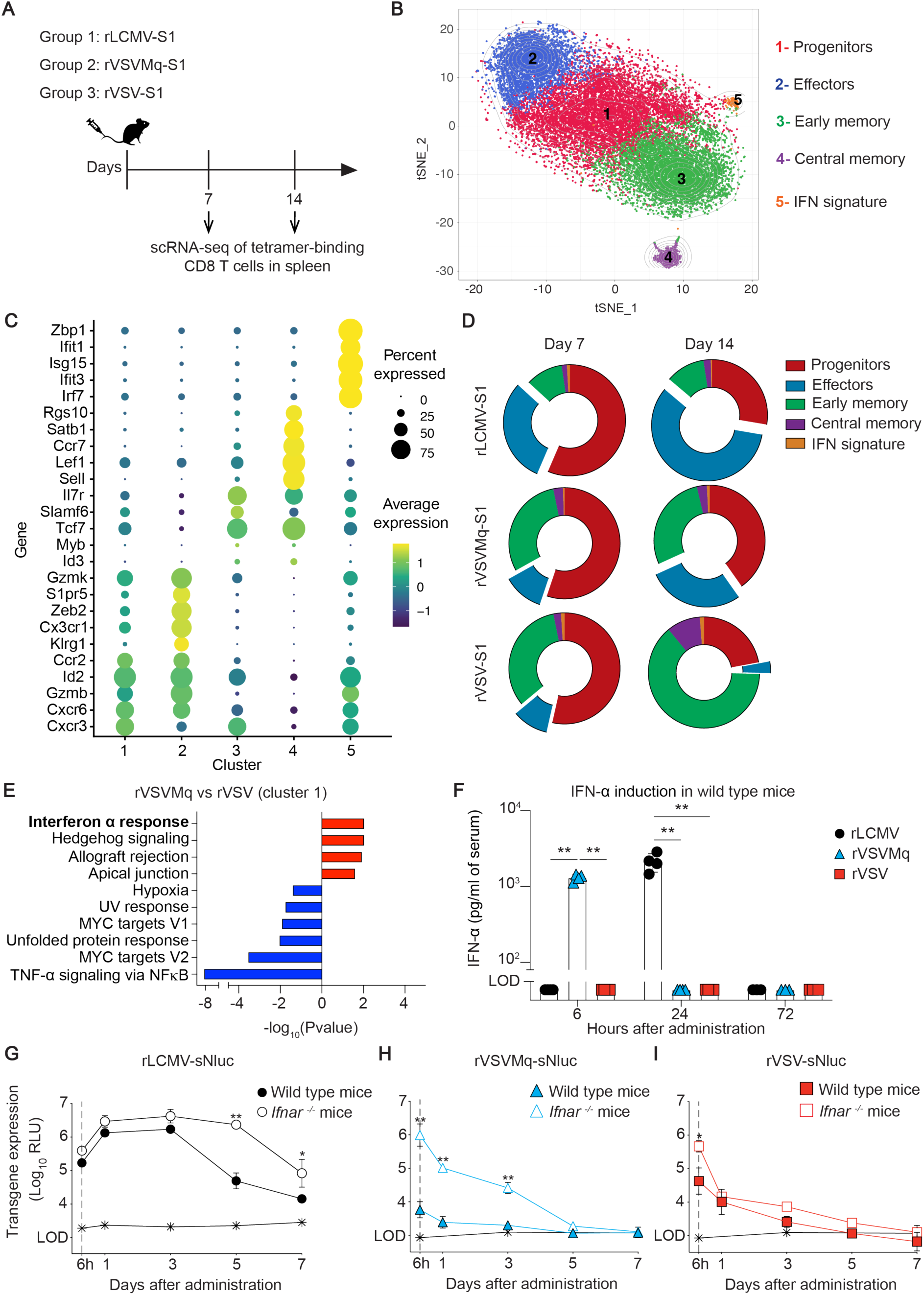
rVSVMq-induced CD8 T cells exhibit transcriptomic signatures of effector differentiation and IFN-I signaling. (A) We immunize WT mice with rLCMV, rVSVMq or rVSV expressing the S1 domain of the SARS-CoV-2 spike protein (Wuhan Hu-1 strain) and spleens were collected at d7 and d14. (B) Clustering of splenic S1 epitope-specific CD8 T cells from two mice per condition (vector used, time point), visualized using t-distributed stochastic neighbor embedding (t-SNE). Each cell is represented by a point and colored by cluster. (C) Dotplot of gene expression levels of selected genes among clusters. (D) Differential abundance of CD8 T cells in each cluster, individually displayed for each vector and time point. (E) Gene set enrichment analysis of cluster 1 cells from day 14, comparing rVSVMq-induced CD8 T cells compared against those induced by rVSV. Gene set enrichment analysis was conducted and gene sets with an FDR < 0.01 are displayed. (F) IFN-α levels in serum of mice immunized with rLCMV-S1, rVSVMq-S1 or rVSV-S1. (G-I) Luciferase activity in serum of WT and *Ifnar ^-/-^* mice immunized with sNluc-expressing rLCMV (G), rVSVMq (H) or rVSV (I). Symbols in (F) represent individual mice (n=4 per group) with bars showing the mean±SEM. Symbols in (G-I) show the mean±SEM of 3 mice per group. One representative experiment of two similar ones is shown in (F, G, H and I). Statistical analyses in (G-I) were performed by two-way ANOVA with Bonferroni’s post-test for multiple comparisons, *: p < 0.05, **: p < 0.01, *p* > 0.05 was considered not statistically significant and is not indicated.

Based on the relative abundance of these T cell clusters we compared the dynamics of CD8 T cell differentiation in rVSV-S1-, rVSVMq-S1- and rLCMV-S1-vaccinated mice (Figs. 3D, S3B). At day seven, all three vectors induced predominantly progenitor cells (cluster 1). T cell populations in the two rVSV-S1-vaccinated groups were similarly distributed, whereas rLCMV-S1-vaccinated animals showed some enrichment in effector cells (cluster 2) and a concomitant reduction in early memory cells (cluster 3). By day 14, the abundance of progenitor cluster 1 cells had proportionally declined in all groups. Of note, however, S1-specific CD8 T cells in rVSV-S1-vaccinated mice were shifted towards the early memory cluster 3, while rVSVMq-S1- and rLCMV-S1-induced responses were enriched in effector-differentiated cluster 2 cells. These transcriptional data corroborated our flow cytometric analyses and documented that rVSVMq-S1-induced CD8 T cell responses were more effector-differentiated, resembling those elicited by rLCMV-S1.

To further compare CD8 T cell responses induced by rVSV-S1 and rVSVMq-S1 and to identify biological pathways enriched in rVSVMq-S1-induced CD8 T cells we performed a gene set enrichment analysis (GSEA). GSEA revealed that rVSVMq-stimulated CD8 T cells from progenitor cluster 1 exhibited significantly higher expression of IFN-α-induced genes (Fig. 3E), suggestive for more pronounced IFN-I signaling upon vaccination with the non-cytolytic rVSVMq-S1 vector than with cytolytic rVSV-S1. To directly test this hypothesis, we assessed systemic IFN-α levels in serum samples collected in the first three days after vector administration. Indeed, rVSVMq-S1 triggered a substantial systemic IFN-α response detectable at six hours after administration, whereas serum IFN-α of rVSV- S1-vaccinated mice remained below detection limits throughout the observation period (Figs. 3F, S3C). In keeping with our earlier report^68^ rLCMV-S1 administration triggered a systemic IFN-α response peaking at 24 hours after immunization.

Higher IFN-I responses to rVSVMq than to rVSV in conjunction with the known role of the VSV M protein as IFN-I antagonist^69,70^ offered a compelling explanation for lower antigen expression levels by rVSVMq than by rVSV (compare Fig. 2B). Hence we analyzed vectored sNluc expression in interferon type I receptor-deficient (*Ifnar*^-/-^) mice (Fig. 3G-I). Both rVSV-sNluc and rVSVMq-sNluc expressed more serum sNluc when administered to *Ifnar*^-/-^ mice instead of WT controls. This difference was, however, much more pronounced for rVSVMq-sNluc than for rVSV-sNluc. At six hours after administration, rVSVMq-sNluc-vaccinated *Ifnar*^-/-^ mice expressed 100-times more sNluc than WT controls (Fig. 3H). These differences persisted over time and by consequence serum sNluc activity in rVSVMq-vaccinated *Ifnar*^-/-^ mice was still over detection limits on day three after administration but had reached technical background levels in WT controls. In contrast, rVSV-vectored sNluc in the serum of *Ifnar*^-/-^ mice was only ∼10-fold higher than in WT mice at six hours after administration, and no statistically significant differences were recorded at later time points (Fig. 3I). rLCMV-vectored sNluc expression levels in *Ifnar*^-/-^ and WT mice were not significantly different in the first three days and persisted for at least seven days in both types of mice, which was supposedly due to the viral nucleoprotein’s ability to dampen IFN-I signaling^71,72^ (Fig. 3G). Collectively, these findings established a correlation between the non-cytolytic replication cycles of rVSVMq and LCMV, the induction of systemic IFN-α responses by these vectors and the sustained effector differentiation of CD8 T cells triggered by the same.

### rVSVMq promotes IFNAR-dependent expansion of antigen-specific CD8 T cells

Virus-induced expansion of CD8 T cells can be promoted by IFN-I, the extent of which depends on the infecting pathogen^73^, prompting us to test whether the same applied to the vectors under study. We administered to mice an IFNAR-blocking antibody or isotype control followed by vaccination with either rVSV-S1, rVSVMq-S1 or rLCMV-S1 expressing S1, and measured S1-specific CD8 T cells over time (Fig. 4A). In rVSV-S1-vaccinated animals IFNAR blockade reduced the frequency of circulating tetramer-binding CD8 T cells in blood 1.6-fold on day seven but failed to exert significant effects at later time points (Fig. 4B). In remarkable contrast, responses to rVSVMq-S1 and rLCMV-S1 were significantly suppressed at all timepoints when IFNAR was blocked. On day thirty, the total number of S1-specific CD8 T cells in the spleen of rVSVMq-S1- and rLCMV-S1-vaccinated mice was 19.7-fold and 5.8-fold reduced, respectively, when IFN-I was blocked, whereas responses to rVSV-S1 were unaffected by the same (Fig. 4C). IFNAR blockade exerted an at least equally pronounced effect on the number of CX3CR1^+^CD27^-^ effector CD8 T cells induced by rVSVMq-S1 and rLCMV-S1 but failed to have a detectable impact on the limited effector CD8 T cell response to rVSV-S1 (Fig. 4D). These data showed that IFNAR signaling was a key determinant of both the magnitude and effector differentiation of CD8 T cell responses to rVSVMq-S1 and rLCMV-S1, but was largely dispensable for rVSV-S1-driven CD8 T cell induction.

**Figure 4.**
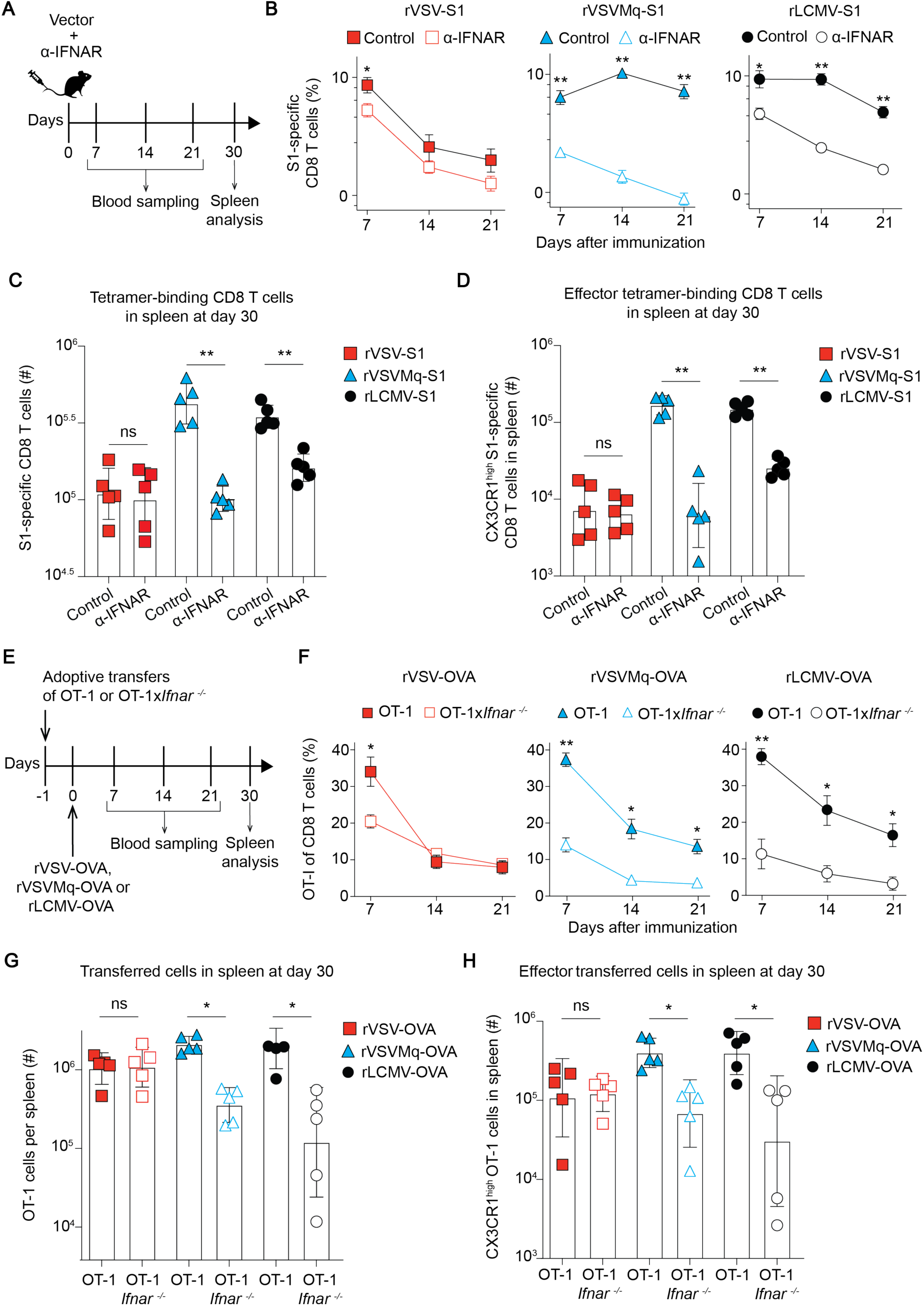
rVSVMq promotes IFNAR-dependent expansion of antigen-specific CD8 T cells. (A) We treated mice with either anti-IFNAR antibody or isotype control and immunized them with rVSV-S1, rVSVMq-S1 or rLCMV-S1 on d0. We analyzed blood over time and collected the spleen on d30. (B) Frequencies of S1 epitope-specific CD8 T cells in blood over time. (C) Splenic count of total S1 epitope- specific CD8 T cells in in spleen on d30. (D) Splenic count of S1 epitope-specific CX3CR1^high^ effector CD8 T cells in spleen on d30. (E) We transferred 2000 OT-1 or OT-1x*Ifnar^-/-^* cells i.v. at d-1 and immunized the recipients at d0 with rVSV-OVA, rVSVMq-OVA or rLCMV-OVA. NK cell-depleting antibody was administered on d-1 and d1. (F) Frequencies of OT-1 and OT-1x*Ifnar^-/-^* cells in blood over time. (G) Total numbers of OT-1 and OT-1x*Ifnar^-/-^* cells in spleen at day 30. (H) Splenic count of OT-1 and OT-1x*Ifnar^-/-^* cells with an effector phenotype (CX3CR1^high^) at day 30. Symbols in (C,D,G,H) represent individual mice with bars showing the mean±SEM. Symbols in (B,F) show the mean±SEM of 5 mice per group. One representative experiment of two similar ones is shown. Statistical analyses were performed with two-way ANOVA with Bonferroni’s post-test for multiple comparisons (B,C,D,F,G,H); ns: not significant; *: p < 0.05, **: p < 0.01.

To determine whether IFN-I-driven CD8 T cell expansion and differentiation depended on CD8 T cell-intrinsic IFNAR signaling, we adoptively transferred OVA-specific OT-1 CD8 T cells that were either *Ifnar*-deficient or -sufficient (OT-1, OT-1x*Ifnar*^-/-^), followed by immunization with rVSV-OVA, rVSVMq-OVA or rLCMV-OVA (Figs. 4E, S4). Recipients were NK cell-depleted to avoid NK cell-mediated killing of *Ifnar*-deficient T cells^74^. On day seven after rVSV-OVA immunization, *Ifnar*-sufficient OT-1 cells reached ∼2-fold higher frequencies than OT-1x*Ifnar*⁻/⁻ cells. This difference was, however, lost by day 14 and both cell populations persisted at comparable frequencies throughout day 21 (Figs. 4F, S4). In contrast, when triggered by rVSVMq-OVA- or rLCMV-OVA, OT-1 cell frequencies in peripheral blood were significantly higher than those of OT-1x*Ifnar*^-/-^ cells from day 7 onwards throughout day 21. Cell counts in the spleen on day thirty paralleled these findings in blood (Fig. 4G). OT-1 and OT-1-IFNAR^-/-^ cell counts were comparable in rVSV-vaccinated animals whereas in rVSVMq- and rLCMV-immunized animals OT-1 cell numbers exceeded those of OT-1x*Ifnar*^-/-^ cell by 5.4- and 8.9- fold, respectively. Analogous findings were made when enumerating CX3CR1^+^CD27^-^ effector memory progeny of OT-1 and OT-1x*Ifnar*^-/-^ cells (Fig. 4H). These results showed that more potent CD8 T cell expansion and effector differentiation in response to rVSVMq-OVA and rLCMV-OVA immunization depended to a significant extent on direct IFN-I signaling to antigen-specific CD8 T cells.

### rVSVMq but not rVSV vaccination offers spatial integration of cognate antigen and IFN-I signals by antigen-specific CD8 T cells

The above experiments indicated that the IFN-I response elicited by rVSVMq was directly sensed by vector-induced CD8 T cells, significantly promoting their expansion and effector differentiation. To further dissect the spatial relationship between vector-transduced cells, IFN-I-producing cells, and antigen-specific CD8 T cells, we adoptively transferred OT-1 cells into wild-type mice, followed by OVA-expressing vector immunization. Spleens were analyzed six, twelve, eighteen and 24 hours later (Figs. 5, S5). IFN-α expressing cells were most abundant at six hours after rVSV-OVA or rVSVMq-OVA immunization and at 24 hours after rLCMV-OVA, corresponding to the latter two vectors’ respective serum IFN-α peak (Fig. S5; compare Fig. 3F). Immunohistochemical analyses of tissue sections revealed that all three vectors targeted predominantly the marginal zone (Fig. 5B-D).

**Figure 5.**
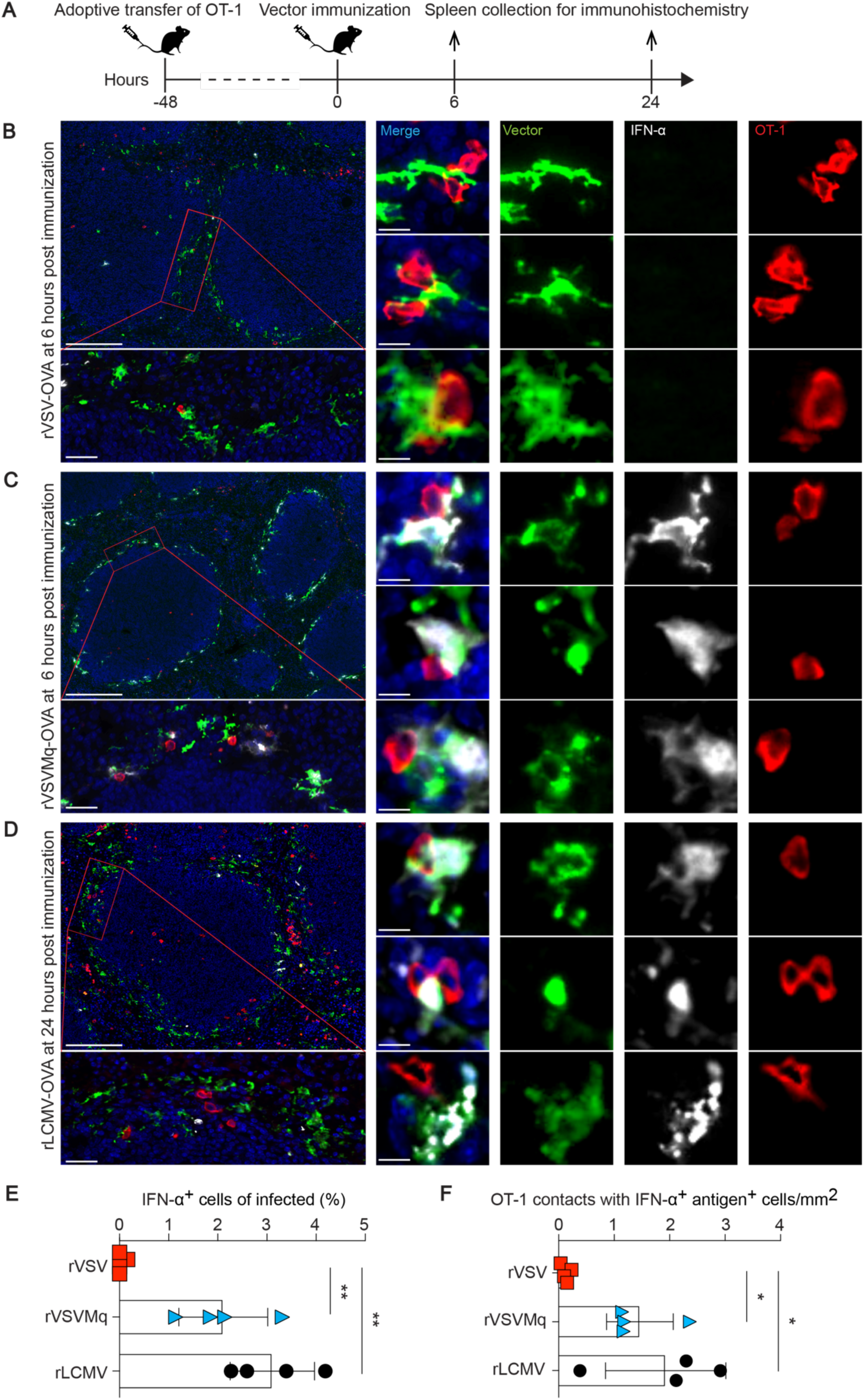
rVSVMq vaccination allows for spatial integration of cognate antigen and IFN-I signals by antigen-specific CD8 T cells. (A) We transferred 2x10^6^ OT-1 cells to mice at -48h and immunized them with either rVSV-EGFP-OVA, rVSVMq-EGFP-OVA or rLCMV-OVA at 0h. Spleens of rVSV-EGFP- OVA (B) and rVSVMq-EGFP-OVA (C) immunized mice were collected at 6h, those of rLCMV-OVA- immunized mice at 24h (D), corresponding to the peak of the serum interferon response in the respective immunization regimens (compare Figs. S5, 3F). (B-C) Spleen sections were stained for cell nucleus (blue), IFN-α (white), CD45.1 (red; OT-1 cells) and GFP (vector infection; green). (D) Spleen sections were stained for cell nucleus (blue), IFN-α (white), CD45.1 (red; OT-1 cells) and LCMV nucleoprotein (vector infection; green). (E) Percentage of IFN-α-expressing cells among virus antigen- positive cells on histological sections. (F) Number of OT-1 cell contacts with IFN-α-positive, virus antigen-positive cells per mm^2^ of histological section. Magnification bars: 100 µm (lowest magnification), 20 µm (intermediate magnification), 5 µm (highest magnification). Representative areas of spleen tissue from whole spleen sections of 5 mice per group, originating from two independent experiments, are shown. Statistical analyses in (E,F) were performed with one-way ANOVA with Tukey’s post-test; **: p < 0.01, *: p < 0.05, *p* > 0.05 was considered not statistically significant and is not indicated.

A substantial proportion of adoptively transferred OT-1 cells were localized in the marginal zone, some of them in immediate proximity to vector-infected cells (Fig. 5B-D), suggesting early cognate interactions. Importantly, in spleens of rVSVMq-OVA- and rLCMV-OVA-vaccinated animals, the marginal zone cells in contact with OT-1 cells often produced IFN-α (Fig. 5C,D). OT-1 cells in the spleen of rVSV-vaccinated animals were in contact with vector-transduced cells, too, but the latter lacked detectable IFN-α production (Fig. 5B). In the spleen of rVSVMq-OVA- and rLCMV-OVA-vaccinated mice, ∼2-3% of vector-infected cells produced IFN-α at levels detectable by histology, while such cells were very rare or absent in rVSV-OVA-immunized animals (Fig. 5E). This differential abundance of IFN-α-producing cells in spleen paralleled serum IFN-α levels in the serum of mice (compare Fig. 3F). Importantly also, OT-1 cells in rVSVMq-OVA and rLCMV-OVA-immunized spleens were often found in contact with vector-infected cells producing IFN-α, while such contacts were virtually undetectable in rVSV-OVA-infected spleens (Fig. 5F).

Taken together, these observations suggested that antigen-specific CD8 T cells engaged with vector-transduced cells in the marginal zone as early as six hours after rVSV-OVA and rVSVMq-OVA immunization and 24 hours after rLCMV-OVA vaccination. Importantly, however, these rVSVMq-OVA- and rLCMV-OVA-infected cells expressed not only cognate antigen but produced also IFN-I, supposedly offering an opportunity for CD8 T cells to integrate T cell receptor and IFN-I receptor signaling.

## Discussion

The induction of robust and durable effector-memory CD8 T cell immunity holds great promise for both prophylactic and therapeutic vaccination against a range of infectious diseases and tumors. Replication-deficient viral vectors represent a preferred modality to induce such responses, and owing to their safety and reactogenicity profile they are attractive platforms for clinical translation. Platform-specific differences in immunogenicity can, however, be substantial, and the biological features determining potent CD8 T cell immunity are mechanistically ill-defined. In this study we demonstrate that rendering a cytolytic vector non-cytolytic can substantially augment its ability to induce durable and protective effector-memory-differentiated CD8 T cell immunity. The pronounced effects of non-cytolytic vector replication were evident at the phenotypic and transcriptional level of antigen-specific CD8 T cells and translated into substantially improved protection against *Listeria monocytogenes* challenge. Mechanistically, we show that non-cytolytic vector replication augments IFN-I production and that IFN-I sensing by CD8 T cells is key for durable effector-memory formation. These observations raise the possibility that IFN-I secretion by antigen-presenting cells in the priming phase of the CD8 T cell response enables the spatio-temporal integration of T cell receptor signals with IFN-I, often referred to as “signal 3” in T cell activation, thereby promoting CD8 T cell proliferation, survival, and effector differentiation^75^.

Earlier studies have shown that IFN-I released upon viral infection can act directly on CD8 T cells to support clonal expansion and memory formation, and in the context of mRNA vaccination IFN-I is essential for CD8 effector T cell differentiation^76^. The relative impact of IFN-I on CD8 T cell responses to infection varied, however, considerably between pathogens^77–81^, and reports on the role of IFN-I in the context of vectored vaccination appear inconsistent. In the context of adenovirus-vectored vaccination IFN-I was shown to limit CD8 T cell expansion^26^. Accordingly, blockade of IFN-I was proposed as a strategy to augment immune responses to various life-attenuated, replicating vaccine vectors, supposedly by increasing and/or extending antigen expression^34^. On the other hand, IFN-I responses elicited by the replication-deficient modified vaccinia virus Ankara (MVA) were required for optimal humoral and cellular immune responses^35,36^. Our study may help reconciling some of these observations by showing that for a specific type of viral vector, rVSV and its variant rVSVMq in our case, the vector’s cytolytic or non-cytolytic replication cycle can dramatically alter the IFN-I-dependance of the CD8 T cell response elicited. Higher systemic IFN-I responses in non-cytolytic infection, potentially due to continuous intracellular replication of non-cytolytic vectors and incessant activation of cytoplasmic RNA sensors such as RIG-I^82^, may simply result in a more pronounced effect on immune responses. Additionally, non-cytolytic infection and higher IFN-I responses may result in altered pathways of innate immune activation, cytokine production^83^ and dendritic cell maturation as reported for VSV expressing non-cytolytic M protein variants^69,84,85^.

As an additional observation of interest, glycoprotein swap between rVSV and rLCMV did not substantially affect primary CD8 T cell responses. Albeit a main determinant of receptor usage and tropism^86,87,88^, vectors pseudotyped with LCMV-GP or VSVG have been reported to infect dendritic cells with comparable efficiency^83^, which may explain the present findings. In stark contrast to prime vaccination, however, the choice of vector glycoprotein had a strong impact on the ability to augment CD8 T cell responses by homologous booster vaccination. The latter was associated with differential vector-neutralizing antibody induction by LCMV-GP- and VSVG-pseudotyped vectors, as previously reported ^46,89,90^.

In summary, this study provides mechanistic insights into the ability of non-cytolytic replication-deficient viral vector systems to induce potent and durable effector-memory CD8 T cell immunity. This refines our understanding how vaccine vector-intrinsic properties are interlinked with adaptive immune responses and helps to rationally harness innate immune signaling for vaccination-induced cellular immunity against intracellular pathogens and tumors.

### Limitations of the study

As evident from the literature^26,34–36,77–81^ the relative importance of IFN-I signals for CD8 T cell responses is context-dependent. Henceforth the relative importance of the present findings may vary somewhat between vector platforms. Moreover, cytokine signals other than IFN-I can exert profound context-dependent effects on CD8 T cell induction as exemplified by the proportionally higher impact of IL-12 on bacterially as opposed to virally induced CD8 T cell responses^91,92^. Last but not least, these findings in mice are yet to be confirmed in humans or non-human primates.

## Materials and Methods

### Animal experiments

C57BL/6 wt mice were originally purchased from Charles River laboratories and bred locally under specific-pathogen-free conditions for colony maintenance and experiments. IFN-α/β-receptor*-* IFN-γ- receptor- *and* RAG1- triple-deficient mice *(Ifnar^-/-^Ifngr^-/-^Rag1^-/-^)*^93^, IFN-α/β-receptor-deficient mice (*Ifnar^−/−^*)^94^ and OT-1 mice^95^, all on a C57BL/6 background, have been described previously. *Ifnar^−/−^* mice and OT-1 mice were intercrossed to obtain OT-1x*Ifnar*^-/-^ mice. Experimental groups were sex- and age-matched. Mice were bred at the ETH Phenomics Center Zurich (EPIC), whereas experiments were performed at the University of Basel in accordance with the Swiss law for animal protection and with permission by the Cantonal Veterinary Office of Basel City. Vector immunizations were performed at a dose of 1E+06 plaque-forming units (PFU) per mouse unless specified differently and administered in a volume of 200 µl into the tail vein.

### Vector generation and titration

rLCMV-OVA has been described^96^. rLCMV-S1 and rLCMV-sNluc were designed and generated analogously, with open reading frames (ORF) encoding the subunit 1 of SARS-CoV-2 spike protein (strain Wuhan-Hu-1, comprising the immunodominant epitope VNFNFNGL) or the secreted nanoluciferase (Promega), respectively, replacing the viral glycoprotein ORF^96^. Unless stated differently, the rLCMV vectors were grown in stably transfected LCMV-GP-expressing BHK-21 (referred to as BHK-23) cells^96^ and were titrated by immunofocus assays^46^ on stably LCMV-GP-expressing 293T cells^96^. For the generation of rVSV and rVSVMq vectors, the VSVG-deleted genomic plasmids pVSVτιG-EGFP and pVSVMqτιG-EGFP^63^ were engineered to express ovalbumin, the subunit 1 of SARS-CoV-2 spike protein or the secreted nanoluciferase genes from an additional viral transcription start – stop cassette. Unless stated differently, rVSV and rVSVMq vectors were grown on VSVG-trans-complementing BHK-G43 cells and titrated as previously described^60^.

In order to generate VSVG-pseudotyped rLCMV vectors, the latter were grown in BHK-G43 cells^63^ to produce rLCMV/VSVG. In brief, expression of VSVG in BHK-G43 cells was induced by adding Mifepristone at 10^-9^ M concentration to the culture medium for 6 hours. After incubation, the cells were infected with rLCMV at MOI: 0.01 for 3 hours, the inoculum was washed away with PBS, and the cells were incubated with fresh media at 37°C. Vectors were harvested 72 hours after infection and stored at -80°C. rVSV vectors were grown in BHK23 cells^96^, infected at an MOI=0.1 for 24h, to produce rVSV/LCMVGP.

### Flow cytometry

Blood samples were collected and stained freshly with antibodies against CD45R/B220 (RA3-6B2), CD8 (53-6.7), CD44 (IM7), CD62L (MEL-14), CD127 (A7R34), Klrg1 (2F1), CX3CR1 (SA011F11), CD27 (LG3A10) and CD43 (1B11) purchased from BioLegend. Subsequently the stained samples were treated with FACS lysing solution (BD Biosciences, Cat. #349202) to remove erythrocytes and fix the cells. For detection of S1-specific CD8 T cells, H2-K^b^ tetramers were conjugated to PE and loaded with the SARS-CoV-2 spike-derived immunodominant peptide epitope VNFNFNGL. For detection of ovalbumin-specific CD8 T cells, H2-D^b^ tetramers were conjugated to PE and loaded with the SIINFEKL epitope. Peptide-MHC tetramers were prepared by the University of Lausanne tetramer core facility. The tetramers were added to the antibody mix for staining. Spleens were mechanically disrupted using a metal mesh and a syringe plunger and cells were counted with an Immunospot S6 device (C.T.L.). For surface staining, splenocytes were incubated with the same cocktail of antibodies and tetramers as described above for blood, with the addition of anti-erythroid cells antibody (TER-119) as a dump gate. Dead cells were stained with Zombie-NIR Fixable Viability Kit (BioLegend, Cat: #423105). Samples were fixed by incubation with 2% paraformaldehyde for 15 minutes at room temperature. All samples were measured on a LSRFortessa flow cytometer (Becton Dickinson) or a 5-laser Aurora spectral flow cytometer (Cytek Biosciences, Fremnont, CA, USA) and analyzed with FlowJo Software (BD Biosciences).

### Virus neutralization assays, determination of serum luciferase activity and ELISA assays for the quantification of S1-specific antibodies and IFN-α in serum

Microtainer tubes (Becton-Dickinson) were used for serum collection. Immunofocus reduction assays were performed for the detection of LCMV neutralizing antibodies^97^. VSV neutralizing antibodies were measured by plaque reduction assays^98^.

Luciferase luminescence in serum samples was measured using the Nano-Glo® Luciferase Assay kit (Promega, Madison, USA), a Saphirell Tecan infinite plex plate reader and white 96-well luciferase plates (Thermo Fisher Scientific Nunc A/S). For determination of the in vivo half-life of sNluc, HEK-293T cells were transfected with the mammalian expression vector pCAGGS expressing the sNluc gene using Lipofectamine 2000 (ThermoFisher). After 24 hours, cell culture supernatant was harvested to determine luciferase activity. Supernatant was administered i.v. into the tail vein of C57BL/6 mice and blood samples were collected over time to determine Luciferase activity. The half-life of sNluc was determined by non-linear regression and one phase decay parameters using GraphPad Prism version 10.2.1.

For detection of S1-specific antibodies, high-binding 96-well flat bottom plates (Sarstedt AG & Co.KG) were coated with 50 ng of spike protein subunit 1 of SARS-CoV-2 (GenScript) per well in 50 µL coating buffer over night at 4 °C. Plates were washed twice with PBS-T (0.05% Tween-20/PBS), then blocked with 200 µL 5% BSA/PBS-T at room temperature for 45 min. In a separate plate, 2-fold serial dilutions of serum samples were performed in blocking solution, and the serially diluted sera were transferred to the spike protein-coated plates and incubated at 37 °C for 1 h followed by five washes with PBS-T. Peroxidase-conjugated polyclonal anti-muse antibody (1:2000 in blocking solution; Jackson, 115-035-062) was added and the plates were incubated at 37 °C for 60 min. After washing five times with PBS-T, HRP activity was detected using ABTS as a chromogen (Pierce) and the absorbance was measured at 405 nm using the Saphirell plate reader (Tecan). Arbitrary units are computed as ln(1000 x A491nm); the limit of detection was determined as the maximum value reached with naive control serum.

Concentrations of IFN-α in mouse serum were determined using the VeriKine Mouse Interferon Alpha ELISA Kit according to the manufacturer’s instructions (PBL Assay Science).

### Listeria challenge

The recombinant Listeria monocytogenes expressing ovalbumin has been described^99^. The bacteria were grown in blood-heart infusion media (Sigma-Aldrich) at 37°C, harvested during the exponential growth phase and washed with phosphate-buffered saline. A dose of 10^3^ colony forming units (CFU) was administered intravenously to mice.

### scRNA-sequencing and bioinformatic analyses

Splenic single cell suspensions were prepared as described for flow cytometric analysis. CD8 T cells were enriched by magnetic-activated cell sorting using the CD8+ T cells Isolation Kit, mouse (STEMCELL), but the antibody mix in the kit was replaced by a mix containing the following biotin-conjugated antibodies from BioLegend: anti-B220 (RA3-6B2), anti-CD19 (6D5), Ly-76 (TER-119), anti- CD4 (H129.19) and anti-CD138 (281-2). Cell suspensions were stained with antibodies against CD45R/B220 (RA3-6B2), CD8 (53-6.7) and DAPI purchased from BioLegend, PE-conjugated H2-K^b^ tetramers loaded with the SARS-CoV-2 spike epitope VNFNFNGL (produced by the University of Lausanne Tetramer core facility). Tetramer-binding CD8 T cells were FACS-sorted (FACS Aria II, BD) and the recovered cells were immediately processed for cell capture and library preparation using the Chromium Next GEM Single Cell 5’ Reagent Kits v2 (Dual Index) with Feature Barcode technology for Cell Surface Protein & Immune Receptor Mapping (CG000330 Rev D) according to the manufacture’s instruction. Paired-end sequencing was performed with the NovaSeq 6000 (Illumina) using an S1 Reagent Kit version 1 (100 cycles) at the Genomics Facility Basel (28 nucleotides for the cell barcode and unique molecular identifier, 8 for the sample index, and 91 for the transcript read).

The mRNA data was mapped to the mm10 genome using the STARsolo framework (STAR version 2.7.10a)^100^. Gene quantification made use of a custom GFT file based on the ensembl gene annotation (version 102). The cell barcode whitelist used for STAR is the file 737K-august-2016.txt provided by 10x Genomics^®^. Other non-standard command line options for STAR included "-- outFilterType BySJout --outFilterMultimapNmax 10 --outSAMmultNmax 1 --outSAMtype BAMSortedByCoordinate --outSAMunmapped Within --soloType CB_UMI_Simple --soloStrand Reverse --outFilterScoreMin 30 --soloCBmatchWLtype 1MM_multi_Nbase_pseudocounts -- soloUMIlen 10 --soloUMIfiltering MultiGeneUMI_CR --soloUMIdedup 1MM_CR --soloCellFilter None -- soloMultiMappers EM --soloBarcodeReadLength 0 --soloCBstart 1 --soloCBlen 16 --soloUMIstart 17 -- soloFeatures Gene --outSAMattributes NH HI AS nM CR CY UR UY GX GN CB UB". Empty droplets were filtered with DropletUtils::emptyDrops (v1.18.1)^101^ with lower threshold set to 100.

The cDNA fastq files for cell surface hashtag data (ADT+HTO) were first trimmed with the tool fastp (version 0.20.1)^102^ and command line options "-b 15 -f 10 -t 0 -A -Q -G -L". After cDNA trimming cell surface hastag data was mapped and quantified with STARsolo, by using the same cell barcode whitelist as for GEX data. The genome index was built from the hashtag sequences as well as the GTF file used for quantification. Additional command line options included "--outSAMtype BAM SortedByCoordinate --outSAMunmapped Within --soloType CB_UMI_Simple --soloStrand Forward -- soloCBmatchWLtype 1MM_multi_Nbase_pseudocounts --soloUMIlen 10 --soloUMIfiltering MultiGeneUMI_CR --soloUMIdedup 1MM_CR --soloCellFilter None --soloBarcodeReadLength 0 -- soloCBstart 1 --soloCBlen 16 --soloUMIstart 17 --soloFeatures Gene". VDJ data were processed for T- cell receptors with 10x Genomics Cell Ranger vdj (v7.0.0). TCR sequencing and gene expression data was intersected and cells were additionally filtere for high confidence TRA and TRB chains. For the 12 samples, non-expressed genes were removed and scaling normalisation was performed using batchelor::multiBatchNorm (v1.18.1)^103^ with mulitplexed sample name as batch factor. The cell cycle phase was assigned with tricycle (v1.10.0)^104^. The top 2000 highly variable genes (HVGs) were selected with scran::modelGeneVar (v1.30.2)^105^ using the multiplexed sample name as blocking variable and not considering B and T cell receptor variable genes. Batch correction on multiplexed sample names was performed with batchelor::fastMNN on 15 neighbours, 50 dimensions and the selected HVGs. For visualisation, t-stochastic neighbour embedding (tSNE) was performed with scater::runTSNE (v1.30.1)^106^ on the batch corrected coordinates.

The cells were clustered on the batch corrected coordinates with scran::buildSNNGraph with 15 neighbours, type "rank" weighting scheme and igraph::cluster_louvain (v2.1.1)^107^ with resolution 0.6. Top markers per cluster were retrieved as high median AUC values obtained from scran::scoreMarkers with blocking factor the multiplexed sample names.

Cluster consisting of mostly cycling cells was removed, and normalisation, dimensionality reduction, batch correction and clustering were repeated as described above. Differential expression analysis was performed per cluster on pseudobulk samples with edgeR (v4.0.16)^102^.

Groups with less than 20 cells were removed, genes were filtered by expression with edgeR::filterByExpr, scaling factors were computed with edgeR::calcNormFactors. The model was fitted with edgeR::glmQLFit and contrasts were tested with edgeR::glmQLFTest. P-values were adjusted for multiple testing with Benjamini-Hochberg (BH) procedure with edgeR::topTags. Gene set enrichment analysis was performed on hallmark collection MH from MSigDb (v2023.2) with limma::camera (3.58.1)^108^ using inter-gene correlation set to 0.01.

### In vivo IFNAR blockade and NK cell depletion

For IFNAR blockade, mice were given 1 mg of anti-IFNAR monoclonal antibody (MAR-1-5A3, BioXcell) i.v. together with viral vectors in the same syringe. Control groups were administered 1 mg of isotype control antibody (MOPC-21, BioXcell) instead. For transfer experiments involving OT-1x*Ifnar*^-/-^ cells, NK cells were depleted by administration of 300 μg of anti-NK.1.1 (PK136, BioXcell) monoclonal antibody to mice on day -1 and on day 1 of vector immunization, as described in^74,109^.

### Adoptive T cell transfer

For OT-1 cell transfer, single cell suspensions were prepared from spleens of naive OT-1 or OT-1x*Ifnar^--^*^/-^ donor mice. Purification of CD8 T cells was performed with magnetic-activated cell sorting (naïve CD8 T+ T cells Isolation Kit, mouse, Miltenyi Biotec). The purity (>90%) was checked before transfer by FACS and then cells were administered i.v. into the tail vein of syngeneic WT C57BL/6J recipients at a dose of 2000 cells/mouse or 2E+06 cells/mouse for the experiments described in Fig. 4 and Fig. 5, respectively. Transferred OT-1 populations expressed the congenic marker CD45.1 for differentiation from the recipient’s endogenous cells.

### Immunohistochemistry

For immunofluorescence analysis of spleen sections, OT-1 cells were MACS-purified (Miltenyi Biotec naive CD8+ T cell isolation kit, mouse) from the spleens of naive donor mice and 2E+06 cells were injected i.v. into the tail vein of wild-type recipient mice followed by vector immunization 2 days later. Animals were sacrificed at the indicated time points and spleens were fixed in 1% PFA in PBS, infiltrated with 30% sucrose and then embedded and frozen in OCT compound (Tissue-Tek, Sakura Finetek Europe). Tissue sections were stained for IFN-α, GFP and CD45.1 as described previously^110^. In brief, cryostat sections were collected on Superfrost Plus Slides (Fisher Scientific), air dried and preincubated with blocking solution (bovine serum albumin with mouse and chicken serum (Sigma) in 0.1% Triton/PBS). Then they were incubated overnight at 4°C with anti-IFN-α antibody (PBL, #32100-1) in 0.1% Triton/PBS. After washing with 0.1% Triton/PBS, anti-rabbit Alexa fluor 647-conjugated (Life, #A31573) secondary antibody was added for 2 hours at room temperature in 0.1% Triton/PBS. After an additional wash, tissue sections were incubated with Dako REAL peroxidase-blocking solution (Dako, K0672) to inactivate endogenous peroxidases and blocked to minimize unspecific binding (PBS supplemented with 2.5% goat serum). Sections were then incubated for one hour with a chicken anti-GFP primary antibody (ICL, #CGFP-45ALY-Z). To visualize the specific signal, anti-chicken HRP-conjugated antibody (Jackson Immuno Research, #103-035-155) followed by amplification with TSA Vivid 520 (Tocris,7534) was used as secondary system. Finally, after a brief blocking step with mouse serum, sections were incubated with a PE-conjugated anti-CD45.1 antibody (Biolegend, #102707) and nuclei were stained with DAPI (Invitrogen, D1306). Slides were mounted in Fluoromount aqueous mounting medium (Sigma-Aldrich, F4680) for imaging. Stained sections were scanned at a resolution of 0.221 μm/pixel using the Panoramic 250 FLASH II Whole Slide Scanner (3DHISTECH). For samples stained for LCMV nucleoprotein, sections were incubated with anti-LCMV-NP antibody (clone VL-4, rat serum) in PBS containing 0.1% Triton X-100 together with the anti-IFN-α antibody.

Images were analyzed in Visiopharm (version 2025.2, Visiopharm, Denmark). For each marker, a U-Net based deep-learning classifier was trained and each cell labelled according to its marker expression profile. Total tissue area, cell counts and per-cell marker positivity were quantified. For cell-cell contact analysis, CD45.1^+^ cell outlines were detected and expanded by 5μm. U-Net based classifiers were then applied to quantify the overlap of vector^+^ IFN-α^+^ and vector^+^ IFN-α^-^ structures with the extended CD45.1 outlines.

### Statistical analysis

Statistical analyses were performed using GraphPad Prism version 9.0 (GraphPad Software). Pairwise comparisons were performed using two-tailed unpaired Student’s t tests. For comparisons across multiple groups, one-way ANOVA followed by Tukey’s post hoc test or two-way ANOVA with Bonferroni’s post-test was applied as reported in the figure legend. Data are reported as mean ± standard error of the mean (SEM), unless otherwise specified. p < 0.05 (*) was considered statistically significant, p < 0.01 (**) as highly significant. Not statistically significant differences (p > 0.05) are not indicated in the figures unless specified.

## Data availability

Raw data of the experimental results reported in this study have been deposited with Zenodo and are publicly available as of the date of publication under the DOI 10.5281/zenodo.17055489. scRNAseq raw data will be deposited with GEO as soon as the platform opens again (presumably once the US government shutdown ends).

## Acknowledgements

We wish to thank Karsten Stauffer for outstanding animal handling and care, Min Lu and Karen Cornille for excellent technical support, Christian Beisel and Mirjam Feldkamp from the Genomics Facility of the University of Basel and D-BSSE of ETH Zurich for single-cell RNA-seq library preparation and next generation RNA sequencing, and Cynthia Saadi for assistance with immunohistochemistry, Morgane Hilpert with the entire DBM flow cytometry core facility for expert cell sorting, sciCORE (http://scicore.unibas.ch/) scientific computing center at University of Basel, and the entire Experimental Virology group for helpful discussions.

## Author contributions

M.C., R.A., A.F.M., T.A.M, D.F., I.W., I.V., D.M., G.Z., and D.D.P. designed the experiments. M.C., R.A, A.F.M., T.A.M, D.F., J.F., F.G., D.B., I.W., I.V., and M.K. conducted the experiments and acquired and analyzed the data. M.C., G.Z., and D.D.P. wrote the manuscript.

