## Supplementary figures for "Non-cytolytic re-engineering of a viral vaccine vector enables durable effector-memory T cell immunity by reinforcing type I IFN induction"

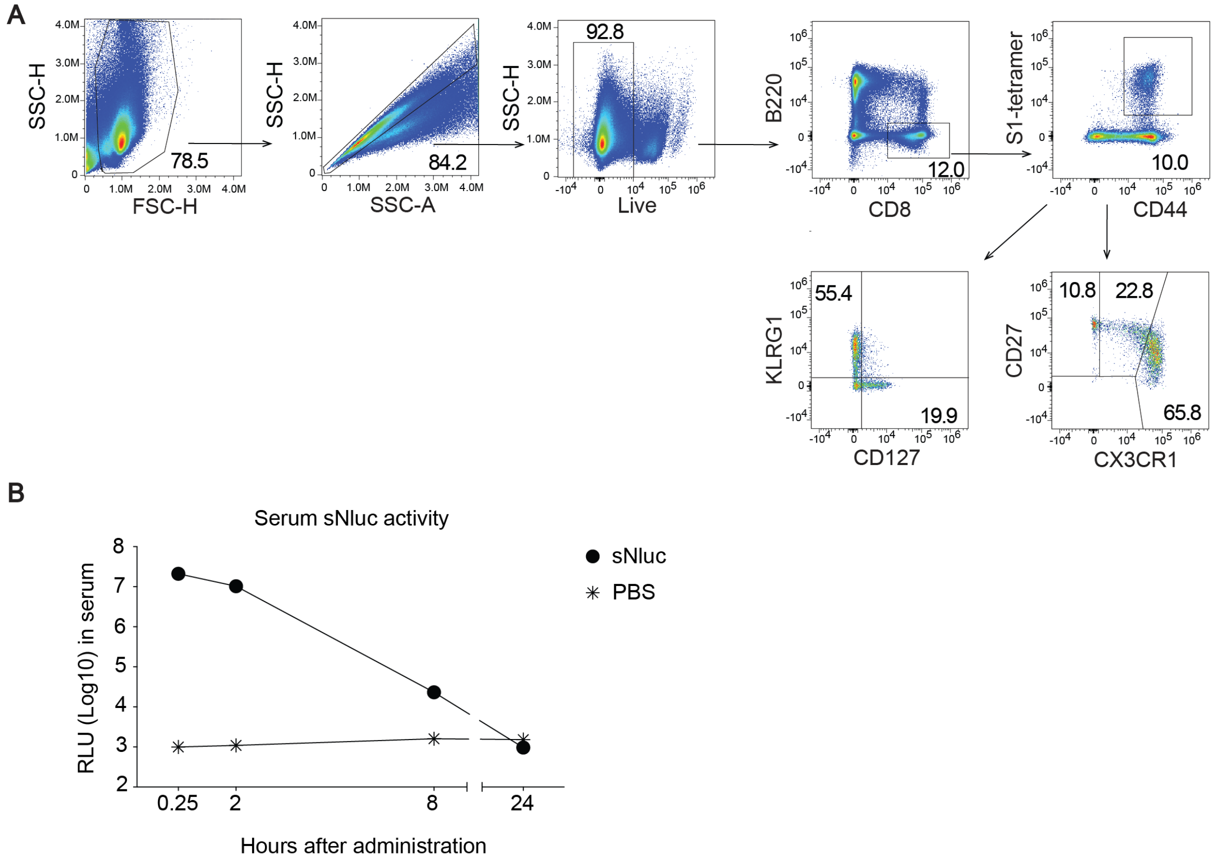


**Figure S1. Gating strategy for epitope-specific CD8 T cells, and *in vivo* half-life of secreted nanoluciferase.** (A) Gating strategy for the enumeration and characterization of tetramer-binding CD8$^{+}$ T cells. (B) Mice were administered cell culture supernatant containing 1E+08 RLU of sNluc or PBS control i.v. and blood was collected to determine luciferase activity in serum over time. Symbols represent the mean±SEM of 4 mice per group. Half-life of sNluc in serum was 0.7 hours, as determined by one-phase decay analysis with non-linear regression using GraphPad Prism.

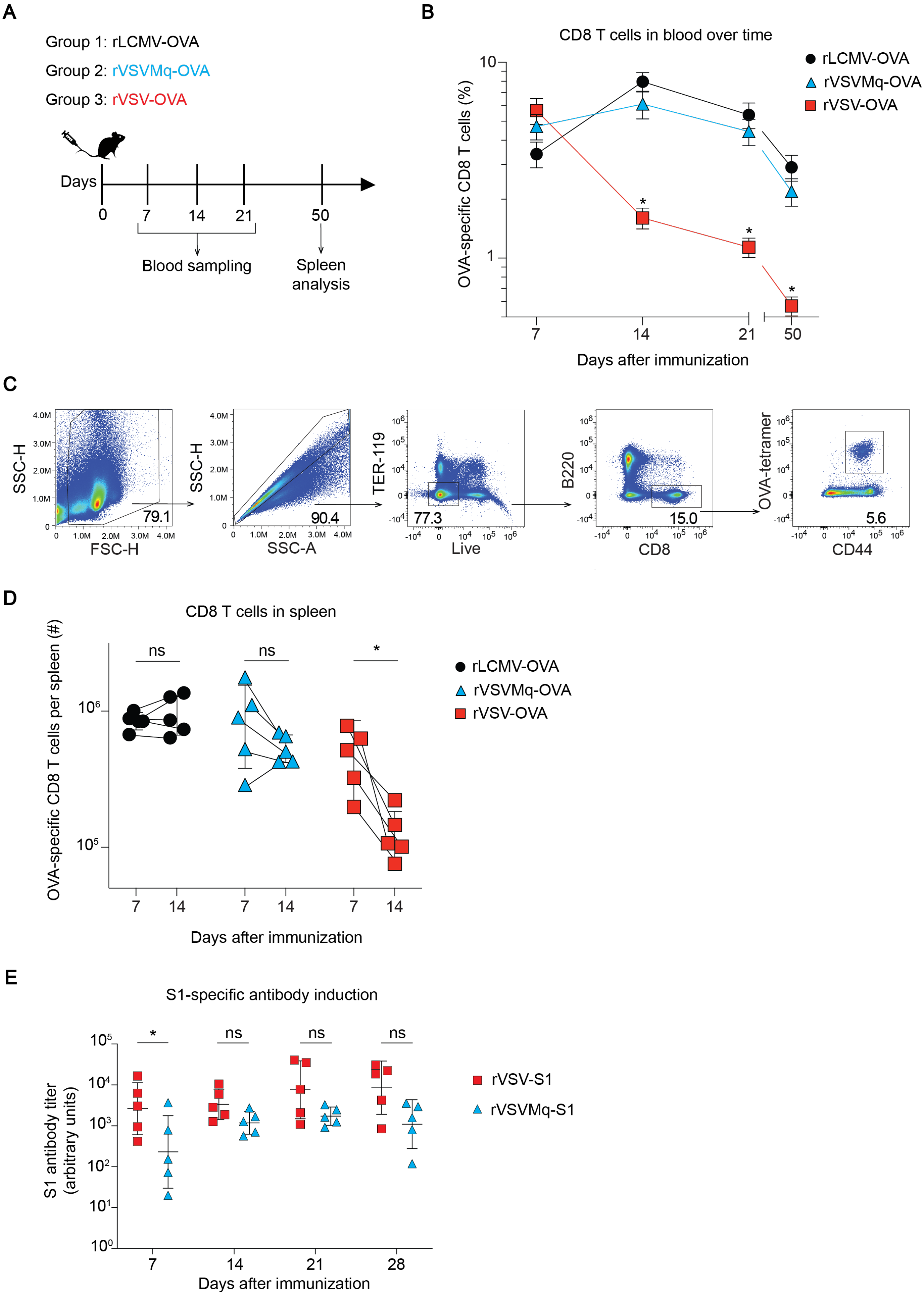


**Figure S2. Kinetics of rVSV-, rVSV-Mq and rLCMV-induced CD8 T cell responses to OVA, and S1-specific antibody induction by rVSV- and rVSV-Mq**. (A-C) We immunized mice i.v. with 1E+05 PFU LCMV-OVA or with 1E+06 PFU of either rVSVMq-OVA or rVSV-OVA on d0, and blood was sampled at several time points. In separate groups of mice, spleens were collected on d7 and d14, respectively. (B) Frequencies of OVA-tetramer-binding CD8 T cells in blood over time. (C) Total numbers of S1-tetramer-binding CD8 T cells in spleen. (E) Mice were immunized i.v. with rVSV-S1 or rVSVMq-S1, and blood was collected at different time points to determine S1-binding serum antibodies by ELISA. Symbols in (B) show the mean±SEM of 5 mice per group, symbols in (D,E) represent individual mice. One representative experiment of two similar ones is shown in (B, D). Statistical analyses were performed by two-way ANOVA with Bonferroni's post-test for multiple comparisons (B,D) or by one-way ANOVA followed by Tukey's post-test (C); **: $p < 0.01$; *: $p < 0.05$; ns.: $p > 0.05$.

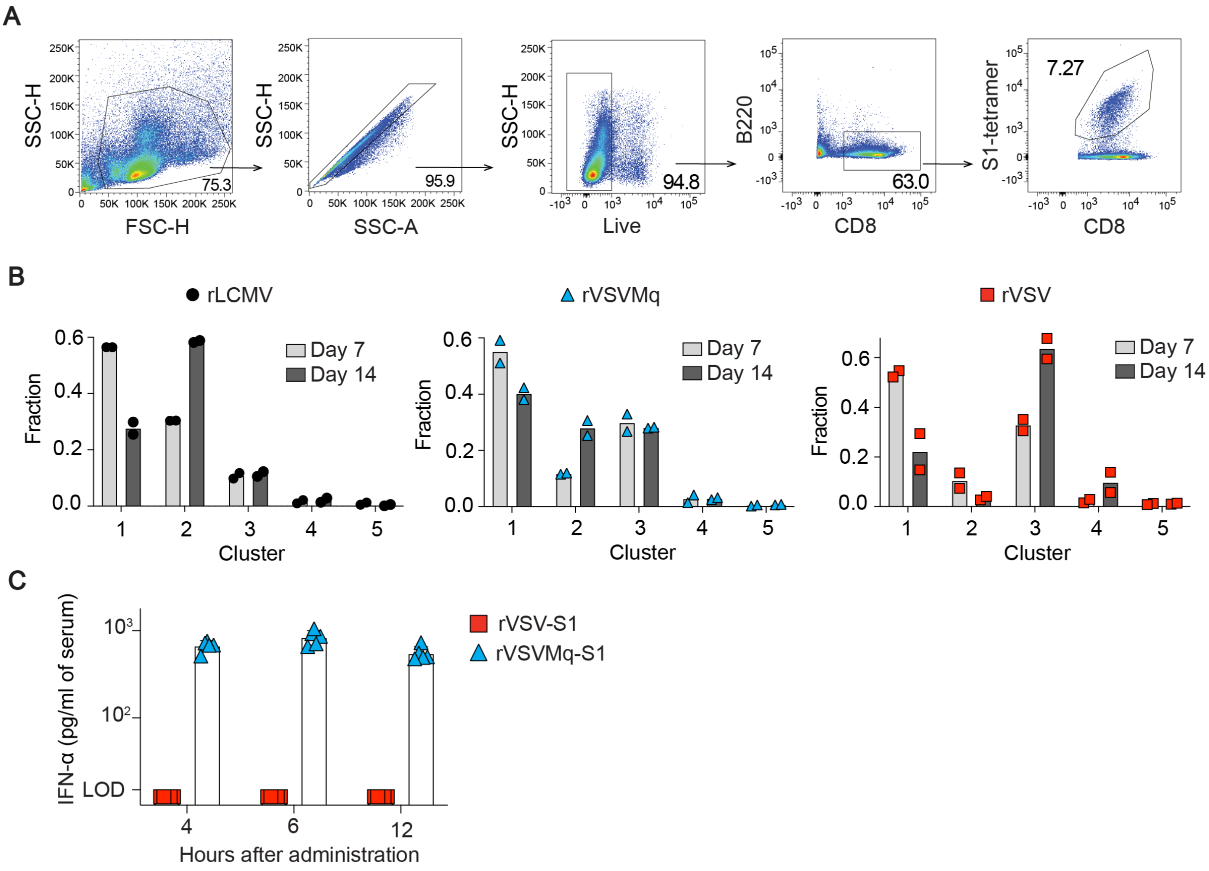


**Figure S3. Gating strategy for FACS-sorting of epitope-specific CD8 T cells, vector-specific distribution of CD8 T cells in clusters, and kinetics of vector-induced IFN-α.** (A) Gating strategy for FACS-sorting of tetramer-binding CD8 T cells to be processed for single cell RNA-sequencing. Prior to sorting, splenic single cell suspensions were enriched for CD8 T cells by magnetic-activated cell sorting. (B) Quantitative assessment of the distribution of rLCMV-S1-, rVSVMq-S1- and rVSV-S1-induced S1-epitope-specific CD8$^{+}$ T cells into clusters on d7 and on d14 of the experiment described in Fig. 3A-E. Symbols show individual mice (n=2 in each group), bars indicate the mean. (C) Time course analysis of IFN-α levels in the serum of mice immunized with rVSV-S1 or rVSVMq-S1. Symbols show individual mice (n=4 in each group), bars indicate the group mean.

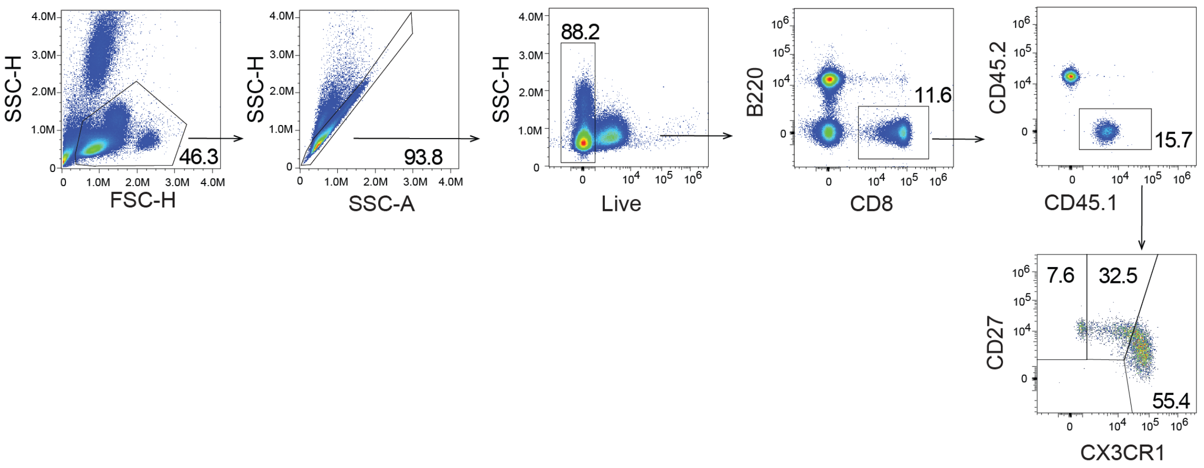


**Figure S4. Gating strategy for the analysis of adoptively transferred OT-1 CD8 T cells.**
Gating strategy for the analysis of adoptively transferred CD45.1$^{+}$ OT-1 and OT-1x*Ifnar*$^{-/-}$ CD8 T cells in CD45.2-congenic recipients.

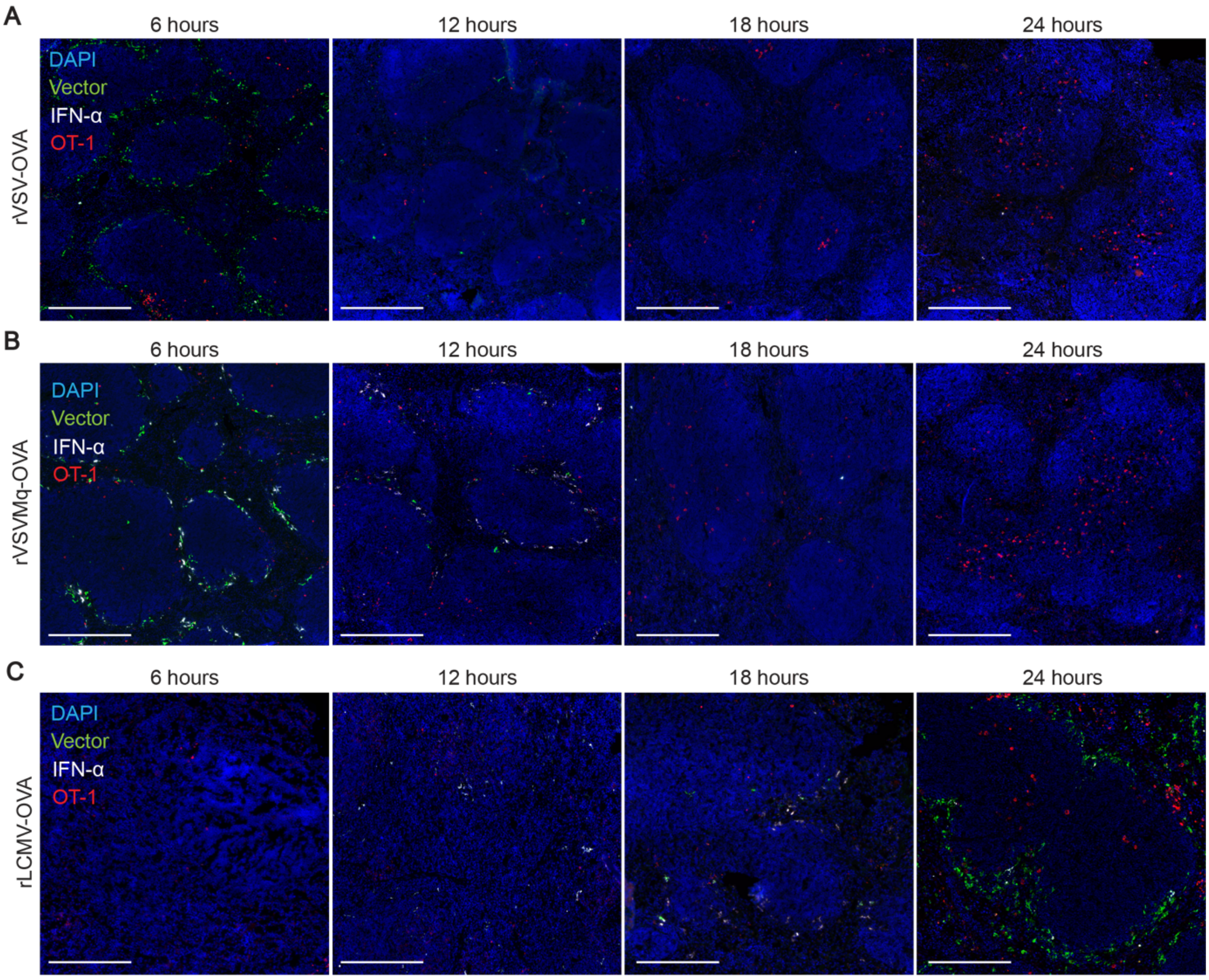


**Figure S5. rVSVMq vaccination allows for spatial integration of cognate antigen and IFN-I signals by antigen-specific CD8 T cells.** We transferred 2E+06 OT-1 cells to mice at -48h and immunized them with either rVSV-EGFP-OVA (A), rVSVMq-EGFP-OVA (B) or rLCMV-OVA (C) at 0h (same experiment as in Fig. 5). Spleens were collected at 6h, 12h, 18h and 24h. In (A,B) spleen sections were stained for cell nucleus (blue), IFN-α (white), CD45.1 (red; OT-1 cells) and GFP (vector infection; green). In (C) spleen sections were stained for cell nucleus (blue), IFN-α (white), CD45.1 (red; OT-1 cells) and LCMV nucleoprotein (vector infection; green). Magnification bars: 100 μm. Representative areas of spleen tissue from whole spleen sections of 4 mice per group from two combined experiments are shown.
